# TSC-Insensitive Rheb Mutations Induce Oncogenic Transformation Through a Combination of Hyperactive mTORC1 Signalling and Metabolic Reprogramming

**DOI:** 10.1101/2020.09.05.284661

**Authors:** Jianling Xie, Stuart P. De Poi, Sean J. Humphrey, Leanne K. Hein, John Bruning, Wenru Pan, Timothy J. Sargeant, Christopher G. Proud

**Author notes:** These authors contributed equally to this work. Corresponding author; mailing address as above.

## Abstract

The mechanistic target of rapamycin complex 1 (mTORC1) is an important regulator of cellular metabolism that is commonly hyperactivated in cancer. Recent cancer genome screens have identified multiple mutations in Ras-homolog enriched in brain (Rheb), the primary activator of mTORC1, that might act as driver oncogenes by causing hyperactivation of mTORC1. Here, we show that a number of recurrently occurring Rheb mutants drive hyperactive mTORC1 signalling through differing levels of insensitivity to the primary inactivator of Rheb, Tuberous Sclerosis Complex.

We show that two activated mutants, Rheb-T23M and E40K, strongly drive increased cell growth, proliferation and anchorage-independent growth resulting in enhanced tumour growth *in vivo*. Proteomic analysis of cells expressing the mutations revealed, surprisingly, that these two mutants promote distinct oncogenic pathways with Rheb-T23M driving metabolic reprogramming and an increased rate of glycolysis, while Rheb-E40K regulates the translation factor eEF2 and autophagy, likely through a differential interaction with AMPK.

Our findings suggest that unique ‘bespoke’ combination therapies may be utilised to treat cancers according to which Rheb mutant they harbour.

## Introduction

The mechanistic target of rapamycin complex 1 (mTORC1) is a serine/threonine kinase that is activated by diverse signals, in particular amino acid availability and growth factors, to drive anabolic processes and inhibit autophagy, a catabolic one^1^. This is achieved through phosphorylation of a range of proteins, the best-characterized of which include the ribosomal protein S6 (rpS6) kinases (S6Ks)^2^ and eukaryotic initiation factor 4E-binding proteins (4E-BPs)^3^. S6Ks phosphorylate rpS6^4^, a component of the 40S ribosomal subunit, and positively regulate cell size, while the 4E-BPs control the initiation of mRNA translation and cell proliferation^4, 5^. 4E-BPs bind to eIF4E in competition with eIF4G to prevent formation of the eIF4F complex thus inhibiting cap-dependent translation^6^. mTORC1-catalysed phosphorylation of the 4E-BPs decreases their affinity for eIF4E allowing eIF4E to form translation initiation complexes and promote translation initiation^7^.

Rheb (Ras homolog enriched in brain) is a small (21 kDa) GTPase that, in its GTP-bound form, activates mTORC1. Hydrolysis of Rheb-GTP is promoted by the GTPase-activating protein (GAP) tuberous sclerosis complex (TSC), comprised of TSC1, TSC2 and TBC1D7, yielding inactive, GDP-bound Rheb^8-10^. The ability of TSC1/2 to impair Rheb function is inhibited by signalling events which are activated by hormones, mitogenic stimuli and growth factors involving the PI3K and MAPK signalling pathways^1, 9, 11^. This involves the phosphorylation of TSC2, the subunit responsible for GAP activity^12^, downstream of oncogenic pathways such as phosphoinositide 3-kinase (PI3K)/protein kinase B (PKB) and rat sarcoma (Ras)/extracellular signal regulated kinase (ERK)^38^. As the pathways upstream of TSC are often aberrantly active in cancer, mTORC1 is also activated in many cancers.

Several mutations in the *RHEB* gene have also been identified in a range of human tumours^13^. Several of these Rheb mutants have been shown to be constitutively active^14-17^. Some mutants are insensitive to TSC’s GAP activity and thus confer constitutive activity on Rheb and mTORC1^16, 18^. In contrast, it has recently been suggested that the constitutively-active Rheb mutant Rheb-Y35N drives continuous mTORC1 activation by activating the MAPK pathway via direct inhibition of AMPKα or heterodimerisation with BRAF^19, 20^. Given that mTORC1 signalling is hyperactive in many cancers and some Rheb mutants can promote oncogenic transformation^15, 19^, it is important to understand the roles of mTORC1 and proteins that control its function in cell function. Here, we show that previously uncharacterised Rheb mutants which occur in human tumours drive constitutive mTORC1 signalling and drive oncogenic transformation and tumour growth *in vivo*. Importantly, in addition to activating mTORC1, some mutants affect other processes, including glycolysis and the translational machinery, effects which differ between mutants. These unexpected findings may require or offer additional therapeutic approaches in treating cancers bearing these mutations.

## Materials and Methods

### Reagents

All reagents were from Merck (NSW, Australia) unless otherwise specified. Bradford assay reagent was from Bio-Rad (NSW, Australia). AZD8055 was purchased from Jomar Life Research (VIC, Australia).

### Cell culture, site-directed mutagenesis and transfection

NIH3T3, HEK293 and HeLa cells were cultured in Dulbecco’s modified Eagle medium (DMEM) containing 10% foetal bovine serum and 1% penicillin/streptomycin at 37°C with 5% (v/v) CO_2_ and regularly tested for mycoplasma contamination. pRK7-FLAG-Rheb vector^21^ was purchased from Addgene and was used to generate Rheb mutants via site-directed mutagenesis. Briefly, mutagenesis primers were used for PCR amplification of pRK7-FLAG-Rheb using pfu Turbo Polymerase (Promega, NSW, Australia) to induce specific mutations. Donor plasmids were digested with *DpnI*. Transfection was performed using Lipofectamine 3000 (Thermofisher Scientific, SA, Australia) according to the manufacturer’s instructions.

### Generation of NIH3T3 cell lines stably expressing Rheb variants

cDNAs encoding wildtype Rheb (Rheb-WT), or Rheb-T23M or E40K were cloned into a pEGFP-N1 vector using the *EcoRI* and *BamHI* restriction sites. Vectors were transfected into NIH3T3 cells which were cultured in the presence of 1 mg/mL G418 for 6 weeks to select stable transfectants. Surviving cells were transferred to 96-well plates at 1 cell/well and monitored for colonies. Rheb expression was determined via SDS-PAGE Western Blot; monoclonal colonies expressing low levels of exogenous Rheb were selected for use.

### SDS-PAGE and immunoblot analysis

SDS-PAGE/Western blot analysis was carried out as previously described ^22^. Briefly, cells were lysed in ice-cold lysis buffer containing 1% Triton X-100, 150 mM NaCl, 20 mM Tris-HCl pH 7.5, 2.5 mM sodium pyrophosphate, 1 mM EDTA, 1 mM EGTA, 1 mM sodium orthovanadate, 1 mM dithiotreitol (DTT) and 1 mM β-glycerophosphate supplemented with protease inhibitor cocktail. Protein concentrations were determined ^23^ and normalized. Equal aliquots of protein (40 μg) were denatured in Laemmli loading buffer, heated at 95°C for 3 min and separated by SDS-PAGE using gels containing 7-13% acrylamide and 0.1%-0.36% bis-acrylamide. Proteins were transferred to nitrocellulose membranes, which were blocked and incubated with primary antibody as indicated (Supplementary Table S1). Fluorescently-conjugated secondary antibody was applied and signals were imaged using a LiCor Odyssey® CLx imager (Millennium Science, VIC, Australia).

### Purification of recombinant GST-Rheb

pGEX-4T2-Rheb vectors were a kind gift from Dr John Blenis (now at Weill Cornell Medicine, New York, NY, USA). Site-directed mutagenesis was performed using pfu Turbo Polymerase. Recombinant GST-Rheb proteins were expressed in and purified from *E. coli* BL21 cells after incubation with 25 µM IPTG (6 h, 30°C). Bacterial lysates were prepared by sonication in ‘Rheb lysis buffer’ (0.2% Triton X-100, 50 mM HEPES-KOH pH 7.4, 140 mM NaCl, 1 mM EDTA pH 8, 1 mM DTT, plus protease inhibitor cocktail). Lysates were incubated with 0.03 U DNase I, 12 mM MgCl_2_ and 0.25 mg/ml lysozyme for 30 min at 4°C, before spinning down at 13,000 x *g* for 10 min at 4°C. Supernatants were incubated with glutathione-agarose (ThermoFisher Scientific) for 2 h at 4°C. GST-Rheb was eluted in 30 mM reduced L-glutathione, 50 mM HEPES-KOH pH7.4, 140 mM NaCl, 2.7 mM KCl, 0.1 mg/ml bovine serum albumin (BSA) plus protease inhibitor cocktail. Eluted GST-Rheb (approximately 1 µg/µl) was then analysed and quantitated by SDS-PAGE and Coomassie staining of resolved proteins and BSA standards.

### GAP Assay

Rheb-GAP assays were performed on complexes immunoprecipitated from HEK293 cells transfected with WT FLAG-TSC1 and WT FLAG-TSC2. Cells were lysed in 1 ml NP-40 lysis buffer (20 mM Tris-HCl pH 7.4, 150 mM NaCl, 1 mM MgCl_2_, 1% Nonidet P-40, 10% glycerol, 1 mM DTT, 50 mM β-glycerophosphate, 50 mM NaF and protease inhibitor cocktail). Flag-tagged TSC1 and TSC2 were then immunoprecipitated with FLAG antibody coupled to protein-G beads for 2 h at 4°C. Immune complexes on beads were washed thrice in IP wash buffer (20 mM HEPES-KOH pH 7.4, 150 mM NaCl, 1 mM EDTA, 1% Nonidet P-40, 1 mM DTT, 50 mM β-glycerophosphate, and 50 mM NaF and protease inhibitor cocktail), and once in 1 ml Rheb exchange buffer (50 mM HEPES-KOH pH 7.4, 1 mM MgCl_2_, 100 mM KCl, 0.1 mg/ml BSA, and 1 mM DTT and protease inhibitor cocktail). GST-Rheb (10 μg) was loaded with 100 μCi [α-^32^P]GTP by incubation for 5 min at 37 °C in 100 μl GTP-loading buffer (50 mM HEPES-KOH pH 7.5, 5 mM EDTA, and 5 mg/ml BSA and protease inhibitor cocktail). After 5 min, 2.5 μl 1 M MgCl_2_, 100 μl ice-cold 50 mM HEPES-KOH pH 7.4, and 20 μl 10 mM GDP were added to the [α-^32^P]GTP-loaded Rheb. GAP assays were initiated by adding 20 μl GTP-loaded Rheb mixture (1 μg GST-Rheb) to each aliquot of FLAG-TSC1/2-protein-G agarose immune complexes. Assays were performed at room temperature with constant agitation for 60 min. Reactions were stopped by adding 300 μl Rheb wash buffer containing 1 mg/ml BSA. Immune complexes were removed by brief centrifugation, and nucleotide-bound GST-Rheb was purified from supernatants by incubating with glutathione beads for 2 h at 4°C. After three washes with Rheb wash-buffer (50 mM HEPES-KOH pH 7.5, 0.5 M NaCl, 0.1% Triton X-100, 5 mM MgCl_2_, 0.005% SDS plus protease inhibitor cocktail), radiolabelled GTP and GDP were eluted from Rheb with 20 µl elution buffer (0.5 mM GDP, 0.5 mM GTP, 5 mM DTT, 5 mM EDTA, and 0.2% SDS) at 68°C for 20 min. One μl of each eluted reaction was resolved by thin-layer chromatography on PEI cellulose with 0.75 M KH_2_PO_4_ pH 3.4 as solvent. Relative levels of [α-^32^P]-labelled GTP and GDP were detected and quantitated with Typhoon phosphor-imager (GE Healthcare, NSW, Australia).

### In silico modelling

*In silico* modelling was carried out using ICM Pro (Molsoft LLC, La Jolla, CA, USA). PDB accession code 6BCU was used as a starting template. Mutants were created in ICM Pro and subjected to 20 rounds of energy minimization and annealing. All molecular visualizations were obtained using the PyMOL graphic tool (the PyMOL molecular graphics system, Version 2.2.3. Schrödinger, LLC).

### 3-(4,5-dimethylthiazol-2yl)-2,5-diphenyltetrazolim bromide (MTT) assay

For MTT assay, 3,000 HEK293 cells expressing Rheb mutants were seeded into 96-well plates. Five mg/ml of MTT was added to each well and plates were incubated at 37°C for 4 h. Medium was carefully aspirated, and crystals were dissolved in 50 µl DMSO. Absorbance was read at 540 nm using a Glomax Discover Multimode Microplate Reader (Promega); cell number was calculated against a standard curve generated by 1:2 serial dilutions of HEK293 cells.

### Bromodeoxyuridine (BrdU) Assay

BrdU Cell Proliferation Assay Kit (Cell Signaling Technology, Danvers, MA #6813) was performed as per the manufacturer’s instructions. Cells were plated at 10,000 cells per well in a 96-well plate 2 h prior to addition of 10 µl of 10x BrdU solution. 2 h after addition of BrdU, cells were fixed/denatured for 30 min. BrdU detection antibody diluted 1:100 and added to cells for 1 h. Cells were washed 3x in wash buffer before addition of HRP-linked secondary antibody for 30 min. Cells were washed 3x before addition of 3,3’,5,5’-tetramethylbenzidine. After 30 min, STOP solution was added and absorbance determined at 450 nm.

### Colony formation assay

To perform colony formation assays, 1 ml base layer containing 0.5% agarose in DMEM was plated in 6-well plates and overlaid with 1 ml of 0.3% agarose in DMEM containing 3,000 NIH3T3 cells stably expressing Rheb-WT, T23M or E40K. Cells were fed every 2-3 days with 1 ml of DMEM containing DMSO, AZD8055 or JAN-384 as indicated. After 6 weeks, cells were stained with 0.05% crystal violet in phosphate-buffered saline containing 2% ethanol. Colonies larger than 100 nm were counted by hand and images captured on a dissecting microscope.

### In vivo tumour model

All animal work was conducted in accordance with the National Health and Medical Research Council’s Care and use of Animals for Scientific Purposes guidelines and with approval (No. SAM339) from the South Australian Health and Medical Research Institute’s animal ethics committee. Tumours were generated by subcutaneous injection of 1 × 10^6^ NIH3T3 cells stably expressing Rheb-WT, T23M or E40K into both flanks of 8-10-week-old, male NOD SCID gamma (NSG) mice (The Jackson Laboratory, Stock number 005557) (5 mice per group generating 10 tumours). Tumours were measured with Vernier callipers and volume calculated (Volume = length x width^2^ x 0.5). When tumours reached a volume of 60 mm^3^, mice were randomly assigned into groups to be injected daily with 20 mg/kg AZD2014 or vehicle (DMSO in PBS containing 5% Tween-80 and 5% PEG) via intraperitoneal injection. After 7 days animals were culled by CO_2_ immersion and tumours were removed.

### Immunohistochemistry

Tumours were embedded in cryo-embedding medium (OCT) and 8 µm sections cut using a Shandon Cryotome E Cryostat at -20°C. Sections were allowed to dry for 30 min before fixation in 10% neutral buffered formaldehyde. Endogenous peroxidase activity was blocked with 3% peroxide and further blocked in 5% BSA to prevent non-specific binding. Indicated primary antibodies were diluted 1:100 in blocking buffer and added to sections at 4°C overnight. Sections were incubated with SignalStain® Boost IHC-Detection Reagent (HRP, Rabbit; Cell Signaling Technologies #8114) at room temperature for 30 min and developed using SignalStain® DAB Substrate Kit (Cell Signaling Technology #8059). Sections were counterstained with hematoxylin for 60 seconds and observed under a light microscope. DAB staining was quantified using ImageJ.

### Mass spectrometry analysis

NIH3T3 cells stably expressing Rheb-WT, T23M or E40K were grown in DMEM either containing or lacking FBS for 72 h. Cells were washed 5x in ice cold PBS, lysed in lysis buffer containing 10% sodium deoxycholate 0.1 M Tris-HCl pH 8.5. Protein concentration was determined using the BCA method and samples normalized. Protein was digested with trypsin (Sigma, T6567). and Lys-C (Wako Chemicals, 129-02541) at a 1:100 ratio of enzyme-to-protein. Peptides were desalted and concentrated using SDB-RPS StageTips as described ^24^ with minor modifications. Briefly, peptides were diluted 1:10 in H_2_O, then 100 µl was mixed 1:1 with loading buffer (99% ethyl acetate, 1% TFA), vortexed vigorously, and then loaded directly onto StageTips packed with 3X layers of SDB-RPS material. Samples were spun through to dryness using a custom 3D printed 96-well StageTip adapter at 1,000 x g. StageTips were washed with a further 100 µl loading buffer, followed by 100 µl wash buffer 1 (99% Isopropanol, 1% TFA), then 100 µl wash buffer 2 (5% acetonitrile, 0.2% TFA). Samples were subsequently eluted directly into clean PCR strip tubes with 60 µl 60% acetonitrile/5% ammonium hydroxide (25% ammonia solution). Samples were then dried in a SpeedVac vacuum concentrator, resuspended in MS loading buffer (2% acetonitrile / 0.3% TFA), and peptides were normalized by A280 absorbance. A Dionex Ultimate 3000 (Thermo Fisher Scientific) UHPLC was connected to a Q Exactive HF-X Orbitrap mass spectrometer, and 1 µg peptides were loaded directly onto a 75 μm I.D., 60 cm column packed with 1.9 μm C18 material (Dr. Maisch ReproSil Pur AQ) and separated over a gradient of 3 to 24% acetonitrile in 0.1% formic acid, over 2 h. Column temperature was maintained at 60°C. The mass spectrometer was operated in data-dependent mode, with one full scan of 350-1,400 m/z performed with resolution 60,000 at a target of 3e6 ions, followed by 20 data-dependent HCD MS/MS scans with a target of 1e5 ions, max IT 28 ms, isolation window 1.4 m/z, normalized collision energy 27, minimum AGC target 1e4, and resolution 15,000. Dynamic exclusion was switched on (30 s). RAW data were processed using MaxQuant version 1.6.6.0, searched against the Mouse UniProt sequence database (June 2019 release) with default settings, with the addition of “Match between runs” (match time window 0.7 min), and “MaxLFQ” enabled. Bioinformatic analysis was performed using Perseus version 1.6.10.43.

### Transwell migration and invasion assays

As previously described ^25^. For migration assays, transwells (8 µm pore size, BD Biosciences, NSW, Australia) were pre-coated with 1% gelatin in serum-free medium, assays were then performed with 1.5 × 10^4^ cells plus 0.5 μg/μl mitomycin C for 24 h using 10 μg/ml collagen and 20% FBS as chemo-attractant. For invasion assays, transwells were pre-coated with Matrigel (1:3 in DMEM, BD Biosciences), assays were performed with 1.5 × 10^4^ cells plus 0.5 μg/μl mitomycin C for 72 h using 10 μg/ml collagen and 20% FBS as chemo-attractants. Transwells were stained with DAPI (4’,6-diamidino-2-phenylindole, 1:20000) and visualized with a Nikon Eclipse Ni microscope (x10 objective lens). DAPI-stained cell numbers were quantified by using the Fiji (Java 8) software.

### Fluorescence Associated Cell Sorting (FACS)

FACS analysis was performed as previously described ^26^. Briefly, tf-LC3 HeLa cells were transfected with the indicated Rheb mutant (Calcium phosphate method) and grown in DMEM containing 10% FBS for 24 h. Cells were collected, strained into a FACS tube and analysed by flow cytometry on a BD LSR Fortessa X20 Analyser (BD Bioscience). Cells were gated on SSC-H and FSC-H. Analysis of red to green fluorescence ratios (mRFP1-H:EGFP-H) was performed using FlowJo v10.6.1 software package.

### Seahorse glycolysis stress test

Agilent Seahorse XF Glycolysis Stress Test (Agilent Technologies, Santa Clara, CA) was performed as per the manufacturer’s instructions. HEK293 cells were transfected with vectors encoding Rheb and allowed to grow for 24 h. 1 × 10^4^ cells were seeded in Seahorse XF microplate and kept in fully-supplemented medium for 6 h to allow attachment. Growth medium was replaced for DMEM lacking FBS overnight. Meanwhile, a 96-well sensor cartridge was hydrated with Seahorse XF calibrant in a non-CO_2_ incubator overnight at 37°C. The following day, sensor cartridge was loaded with 10 mM glucose in port A, 1 µM oligomycin in port B and 50 mM 2-deoxyglucose in port C. Cells were prepared for the assay by replacing growth medium with Seahorse XF Base Medium supplemented with 2mM glutamine, and adjusted to pH 7.4 for 1 h. Glycolysis stress test was then conducted in a Seahorse XFe96 Analyser using the Wave software package. Extracellular acidification rate (ECAR) was normalized to Hoechst stain and graphed using GraphPad Prism 8.

## Results

### Rheb mutants commonly found in cancer promote hyperactive mTORC1 signalling independently of MAPK

To assess the extent of Rheb mutations in cancer, we compiled cancer genomic screen data from three databases of patient-derived cancer mutations [COSMIC, The Cancer Genome Atlas and the Broad Institute Cancer Cell Line Encyclopedia (Supplementary Table S3)]. We found that, similarly to Ras mutations, Rheb mutations occur in a wide variety of cancers (Fig. 1a). We identified several mutations that were present in multiple samples with the four most common mutations being T23M, Y35N, E40K and Q57* (Fig. 1b/c).

**Figure 1.**
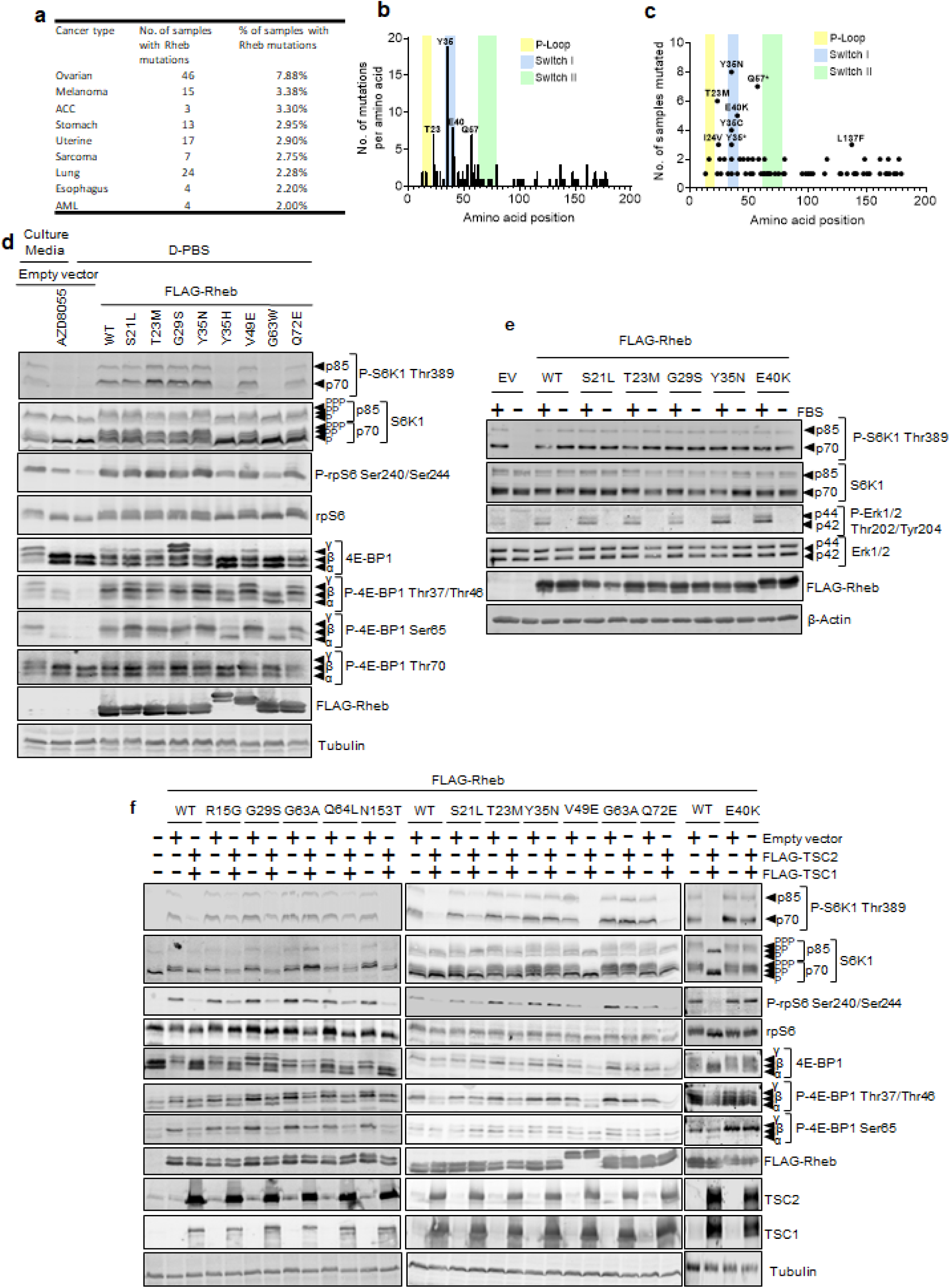

To determine whether these mutations drive hyperactive mTORC1 signalling, we employed site-directed mutagenesis to create selected mutations in a vector encoding FLAG-tagged Rheb. We then transfected HEK293 cells with these vectors, one encoding Rheb-WT or an empty vector, as negative control. To assess the effect of the Rheb mutants on mTORC1 activity, transfected cells were cultured for 16 h in DMEM lacking FBS, conditions under which mTORC1 is generally inactivated, while active Rheb can restore mTORC1 signalling^27^. Cell lysates were then analysed by immunoblot for downstream effectors of mTORC1, including its direct substrates 4E-BP1 and S6K1, and the S6K1 substrate rpS6.

As expected, in fully supplemented growth medium, conditions where the TSC complex does not act on Rheb-GTP, WT Rheb and mutants did not promote levels of mTORC1 signalling above those seen in cells transfected with empty vector (Supplementary Fig. S1a). As expected, in every case, the mTOR kinase inhibitor AZD8055 inhibited the phosphorylation both of S6K1 at Thr389 and of rpS6 at Ser240/Ser244 (Supplementary Fig. S1a) in fully supplemented medium showing the effect of every Rheb variant tested is mediated through mTOR. In contrast, when expressed at similar levels in cells that were subsequently deprived of serum (conditions where mTORC1 signalling is normally inactivated), wild-type Rheb and certain Rheb mutants supported the sustained phosphorylation of S6K1, rpS6 and 4E-BP1 (Fig. 1d). This indicates that those mutants are active, but does not differentiate clearly between their abilities to activate mTORC1 signalling. We noted that the Rheb mutants Rheb-Y35H, -V49E, and -E40K showed altered mobility on denaturing polyacrylamide gel electrophoresis. To try to determine the cause, we performed mass spectrometry to compare post translational modifications of Rheb-WT and E40K; however, we were unable to detect any differences that would explain the observed band shift (data not shown)

It has previously been reported^19, 20^ that Rheb-Y35N drives hyperactive mTORC1 signalling through constitutive activation of the MAPK pathway. We also noted that some Rheb mutants cause greater activation of ERK signalling (P-Erk1/2) than WT Rheb (Supplementary Figure S1e), suggesting this might contribute to their abilities to activate mTORC1. However, P-Erk-1/2 was abolished when cells were deprived of serum which suggests that Rheb mutants do not promote truly constitutive MAPK signalling under the conditions tested here (Fig. 1e; quantified in Supplementary Figure S1b). Furthermore, treatment with the MEK inhibitor AZD6244 did not decrease mTORC1 signalling in cells expressing Rheb mutants (Supplementary Fig. S1c), suggesting that constitutive mTORC1 activation is promoted independently of the MAPK pathway.

Transferring cells to a medium lacking amino acids, such as D-PBS, causes a profound inhibition of mTORC1 signalling. When we tested WT and selected Rheb mutants under these conditions, it was clear that, while WT Rheb promotes mTORC1 signalling, several mutants (T23M, G29S, Y35N) do so more strongly (strongly suggest we move these data into main Fig. 1 and quantify them). Thus, under these conditions, it appears that some Rheb mutants are more potent activators of mTORC1 signalling than WT Rheb.

### Rheb are insensitive to inhibition by TSC

It was possible that the ability of some ectopically-expressed Rheb mutants to promote mTORC1 signalling under starved condition might arise because their levels (likely well above endogenous Rheb) exceeded those of TSC1/2, rather than due to intrinsic constitutive activity. To test this, we co-expressed Rheb mutants alongside vectors encoding FLAG-TSC1/2, to increase the available levels of this complex (Fig. 1f). Strikingly, TSC1/2 decreased phosphorylated P-S6K1 Thr389 or P-rpS6 Ser240/244 in cells expressing wild-type Rheb but not in cells expressing Rheb-G63A. Likewise, phosphorylation of these proteins was less sensitive to TSC1/2 in cells expressing Rheb-T23M, -Y35N or -E40K than in cells expressing Rheb-WT. Cells expressing Rheb-S21L, -G29S or -Q64L still displayed increased P-S6K1 levels compared to cells expressing Rheb-WT when co-expressed with TSC1/TSC2; however, P-S6K1 was more sensitive to TSC1/2 in cells expressing Rheb-S21L, G29S or Q64L than in those expressing Rheb-T23M, Y35N, E40K or G63A (Fig 1e; quantified in Supplementary Fig. S1d). These data suggest that certain Rheb mutants promote hyperactivation of mTORC1 because they are insensitive to TSC1/2..

To assess whether specific Rheb mutations affected the ability of TSC1/TSC2 to promote the hydrolysis of Rheb-bound GTP, we performed GAP assays using recombinant GST-tagged Rheb purified from *E. coli* BL21 cells and FLAG-tagged TSC1/TSC2 co-expressed in, and immunoprecipitated from, HEK293 cells (Fig. 2a). To validate the assay, Rheb-WT and Rheb-Y35N were ‘preloaded’ with [α-^32^P]GTP and then incubated with or without TSC1/TSC2. TSC1/TSC2 efficiently promoted hydrolysis of [α-^32^P]GTP bound to Rheb-WT, but not GTP bound to Rheb-Y35N (Fig. 2b), which has been previously shown to be resistant to TSC1/2’s GAP function^19, 28^. This validated the Rheb-GAP assay, allowing us to test further mutants. TSC1/TSC2 promoted hydrolysis of [α-^32^P]GTP bound to Rheb-T23K, Y35C and N153T similarly to Rheb-WT; in contrast, hydrolysis of GTP bound to Rheb-R15G and G63A was partially resistant to stimulation by TSC1/2. Furthermore, TSC1/TSC2 complexes were completely unable to promote hydrolysis of [α-^32^P]GTP bound to Rheb-S21L, T23M, G29S, Y35N or E40K. This striking difference is notable as this property would allow these Rheb mutants to escape inhibition by the TSC1/TSC2 tumour suppressor complex (Fig. 2c/d). These data suggest that the Rheb mutants S21L, T23M, G29S, Y35N and E40K promote hyperactive mTORC1 signalling (under serum-starved conditions, Fig. 1d) because they are insensitive to the GAP function of TSC.

**Figure 2.**
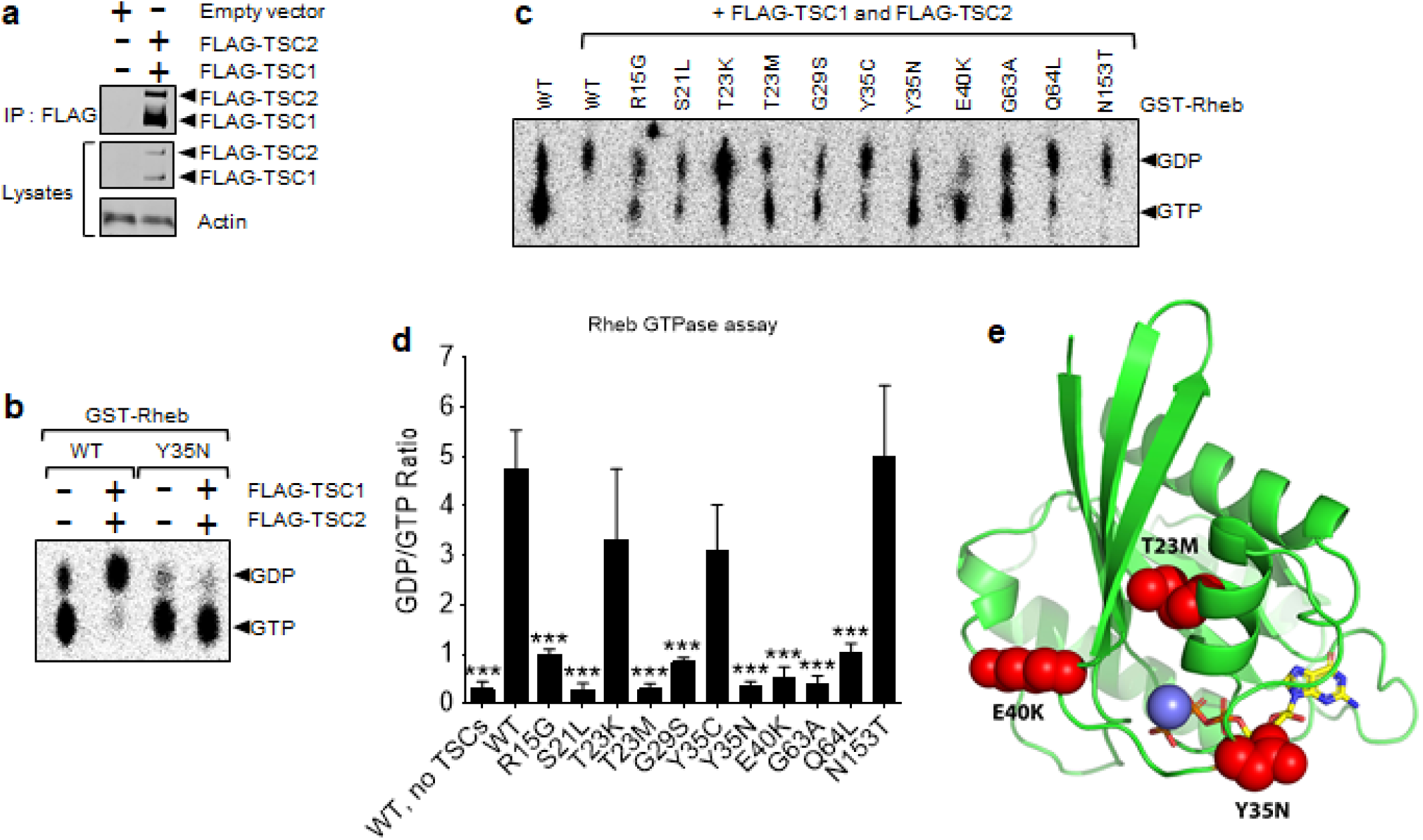

To try to gain further insights into the molecular basis of these differences, we performed *in silico* modelling for Rheb-T23M, -Y35N and -E40K. A loop covers the active site (Fig. 2e), implying importance in substrate binding due to its mobility and direct contacts with substrate. Additionally, there is an α-helix which flanks the substrate, GTP, making direct contacts with the bound GTP. The T23M side chain lies between the afore-mentioned α-helix and the β-sheet (Fig. 2e). The larger Met here likely forces the α-helix closer to the substrate, potentially altering binding of the guanine nucleotide and affecting catalysis. E40K is at the base of the loop that covers the active site (Fig. 2e); this mutation would likely perturb the loop’s conformation, altering its mobility and substrate binding, hydrolysis and/or release.

### Rheb T23M and E40K drive an increase in cell growth, proliferation, and anchorage independent growth

To determine whether Rheb mutants can drive cell proliferation and increased total cell mass, HEK293 cells were transfected with vectors encoding the Rheb mutants Rheb-S21L, T23M, G29S, Y35N and E40K and subjected to MTT and BrdU incorporation assays with/without FBS. MTT assays, which estimate cell number (and thus, over time, cell proliferation) by measuring mitochondrial activity^29^, were performed every 24 h for 7 days to generate growth curves. While there was no difference in cell number between cells expressing WT Rheb or its mutants in fully-supplemented medium (Supplementary Fig. S2a), we saw striking differences when cells were placed in 0% FBS(Fig. 3). While there was no increase in cell number in cells expressing Rheb-WT, each of the Rheb mutants promoted increased cell number (Fig. 3a). In particular, cells expressing Rheb-T23M grew to a higher density than any other mutants, while Rheb-Y35N and E40K showed similar effects on cell number, which were greater than those of Rheb-S21L and G29S (or Rheb-WT).

**Figure 3.**
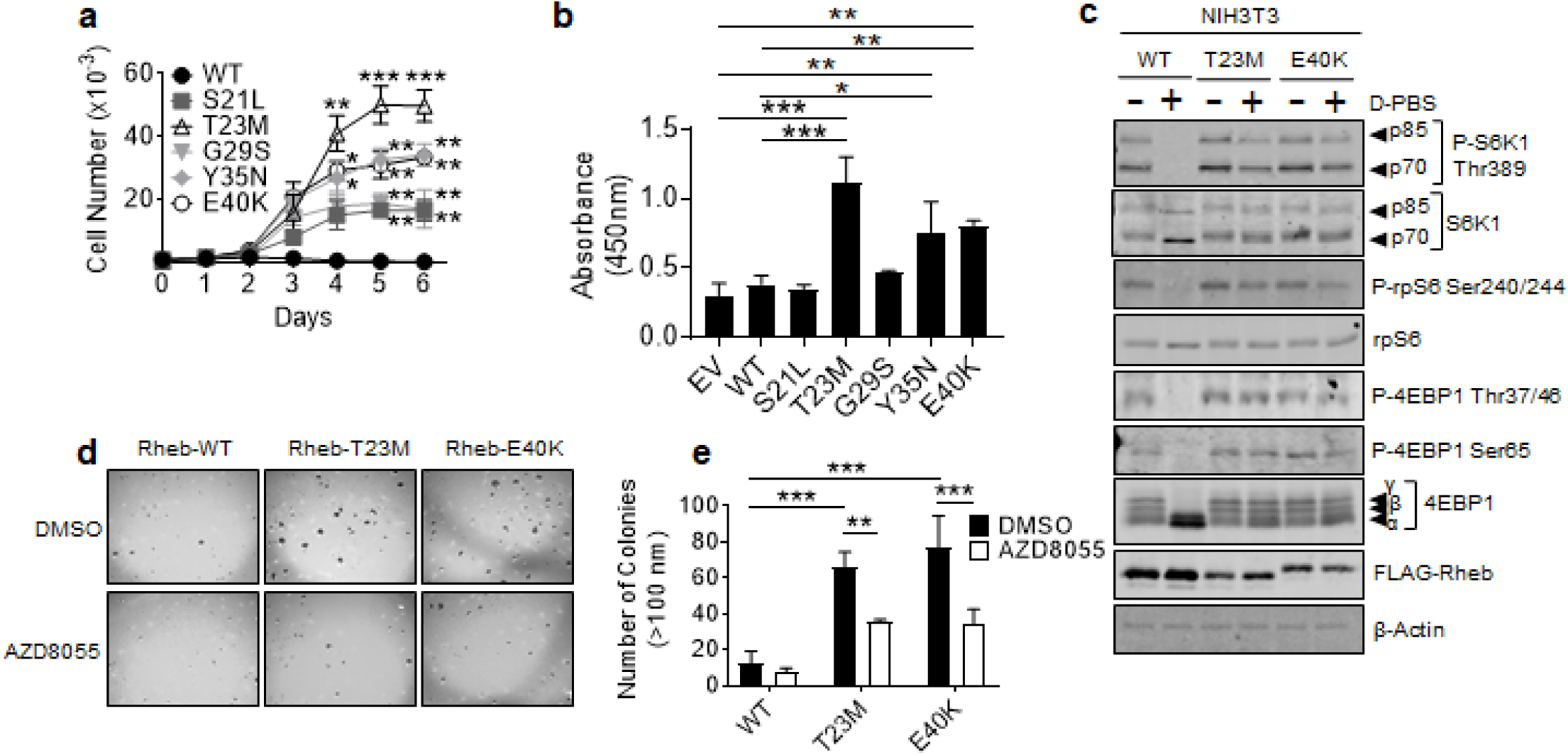

However, as the potential of a cell to reduce MTT depends on its metabolic rate, it is an imprecise measure of cell number and proliferation. As it was therefore important to measure cell proliferation by other means, we performed BrdU incorporation assays, which measure DNA replication. Cells expressing Rheb-T23M, Y35N and E40K cells showed significantly greater BrdU incorporation than those transfected with Rheb-WT or the empty vector, either in the absence or presence of FBS in the growth medium (Fig. 3b; Supplementary Fig. S2b).Rheb-S21L and G29S caused no increase in BrdU incorporation in either the presence of absence of FBS (Fig. 3b; Supplementary Fig. S2b). Taken together, these data strongly suggest that Rheb-T23M, Y35N and E40K each increase cell proliferation under serum-starved conditions. Given that Rheb-Y35N has previously been characterised^19, 20, 28, 30^, and Rheb-S21L and G29S do not increase cell proliferation, we subsequently focused further on Rheb T23M and E40K.

As hyperactive mTORC1 signalling occurs in multiple cancers^31^, we hypothesised that Rheb mutants that drive hyperactive mTORC1 signalling may be oncogenic and drive tumour growth. To test this hypothesis, we stably expressed Rheb-WT, T23M or E40K in mouse NIH3T3 cells (Fig. 3c; Supplementary Fig. S3a). To determine whether Rheb mutants cause oncogenic transformation of NIH3T3 cells, we performed assays for anchorage-independent growth in soft agar. Cells expressing Rheb-T23M and E40K grew robustly in soft agar, forming large colonies, while cells expressing Rheb-WT did not (Fig. 3d; quantified Fig. 3e). Treatment with AZD8055 prevented this, indicating that Rheb-T23M and -E40K each promote oncogenic transformation in a manner dependent upon mTOR signalling, a pathway that is accepted as being tightly associated with cancer development *in vivo*^32^.

### Rheb-T23M and E40K are oncogenic and drive increased tumour growth in vivo

To further test whether Rheb mutants drive tumour growth, NIH3T3 cells stably expressing Rheb-WT, T23M or E40K were injected subcutaneously into the flanks of 8-10-week old, male, immunodeficient NSG mice. When tumour volumes exceeded 60 mm^3^, some animals were treated with 20 mg/kg AZD2014 (an mTOR inhibitor suitable for use *in vivo*33) or vehicle via intraperitoneal injection daily for 7 days. Tumours from cells expressing Rheb-T23M and E40K grew significantly faster than those expressing Rheb-WT so that, over the time of the experiment, Rheb-T23M and E40K tumours also reached a greater volume (approximately 1,000 mm^3^; the permitted end-point) and mass (approximately 1,200 mg) compared to Rheb-WT tumours (approximately 300 mm^3^ and 250 mg). Treatment of mice with AZD2014 greatly impaired tumour growth indicating that Rheb-driven tumour growth depends on mTOR (Fig. 4a-c). Interestingly, while rates of growth and tumour mass were similar between Rheb-T23M and E40K-generated tumours, the times for tumour growth to become evident itself differed, with Rheb-E40K-driven tumours appearing on average 5 days earlier than Rheb-T23M or Rheb-WT tumours (Supplementary Fig. S3b). Immunohistochemical staining with P-rpS6 Ser240/244 and P-ERK Thr202/Tyr204 antibodies was performed on 8 µm sections to assess the mTORC1 and MAPK pathways within the tumours. As expected, tumours expressing Rheb-T23M and E40K showed increased P-rpS6 compared Rheb-WT tumours (Fig. 4d; quantified Supplementary Fig. S3c) while P-rpS6 was absent from tumours from animals treated with AZD2014. There was no difference in P-ERK between any of the tumours showing that MAPK was not activated (Fig. 4d; Supplementary Fig. S3d).

**Figure 4.**
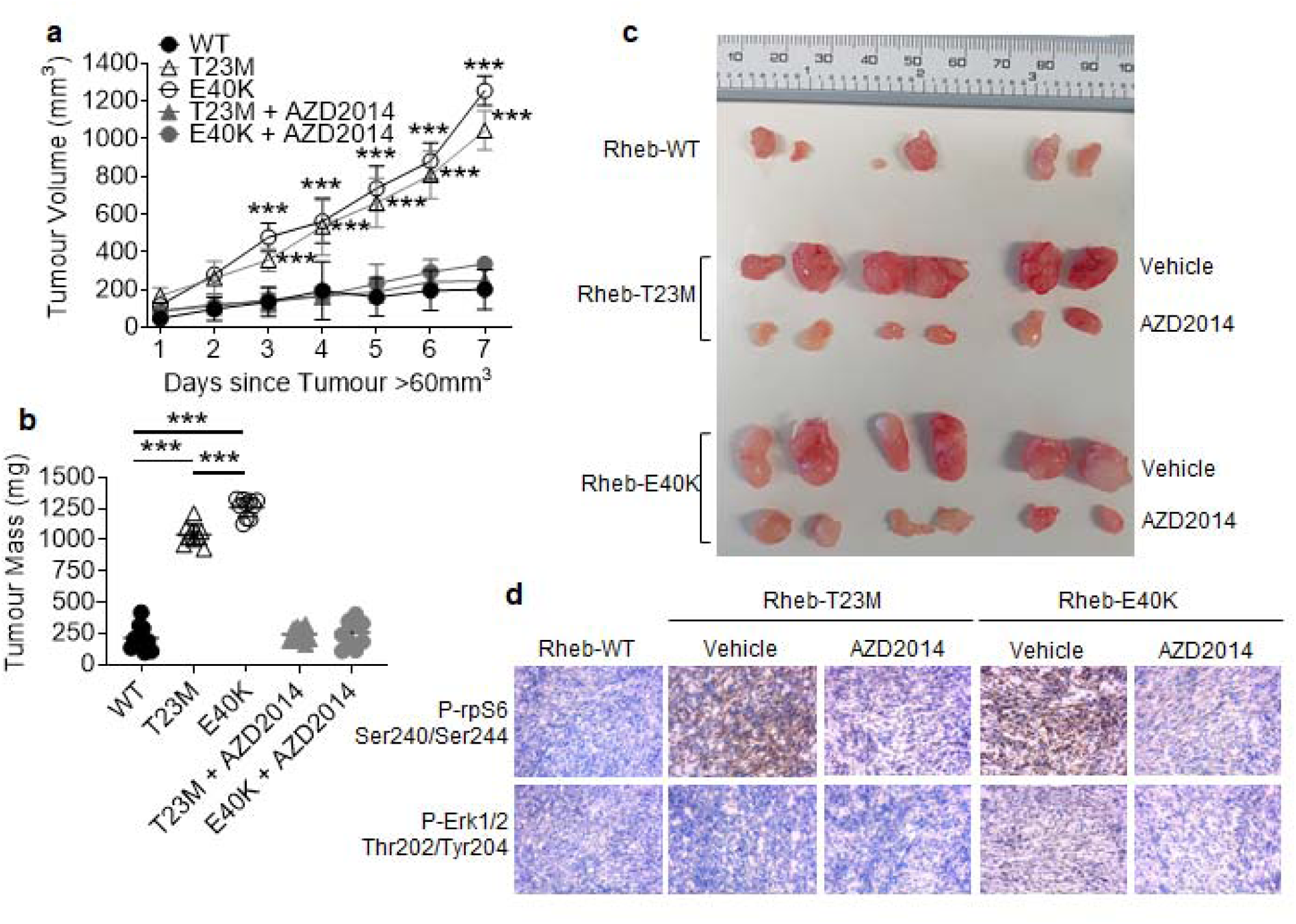

### Rheb-T23M and E40K promote cancer through distinct changes to the proteome

Given we observe hyperactive mTORC1 activation driven by four previously-unknown Rheb mutants, but only two (Rheb-T23M and E40K) appear to be oncogenic while Rheb-S21L and G29S do not, we hypothesised that Rheb-T23M and E40K may exert additional, perhaps mTORC1-independent, functions that aid tumour growth. To test this, we starved NIH3T3 cells stably expressing Rheb-WT, T23M and E40K of serum for 72 h before performing HPLC-mass spectrometry (MS) to evaluate changes in the global proteome (Fig. 5a; Supplementary Fig. S4a-c); we adopted this approach as serum-starvation inactivates WT Rheb and downstream mTORC1 signalling, whereas the mutants remain active (Fig. 1d). Unexpectedly, we found that Rheb-T23M and E40K induce distinct changes in the proteome (Fig. 5b) with Rheb-T23M appearing more similar to Rheb-WT than Rheb-E40K. In cells expressing Rheb-T23M, we observed increases in the key glycolytic enzyme pyruvate kinase (PKM) In contrast, we found that Rheb-E40K upregulated proteins involved in protein synthesis or its control, such as La Ribonucleoprotein Domain 4 (LARP4). We also found that integrin signalling proteins were either increased or decreased consistent with their role in either activating or inhibiting cell migration by both Rheb-T23M and E40K (Fig. 5).

**Figure 5.**
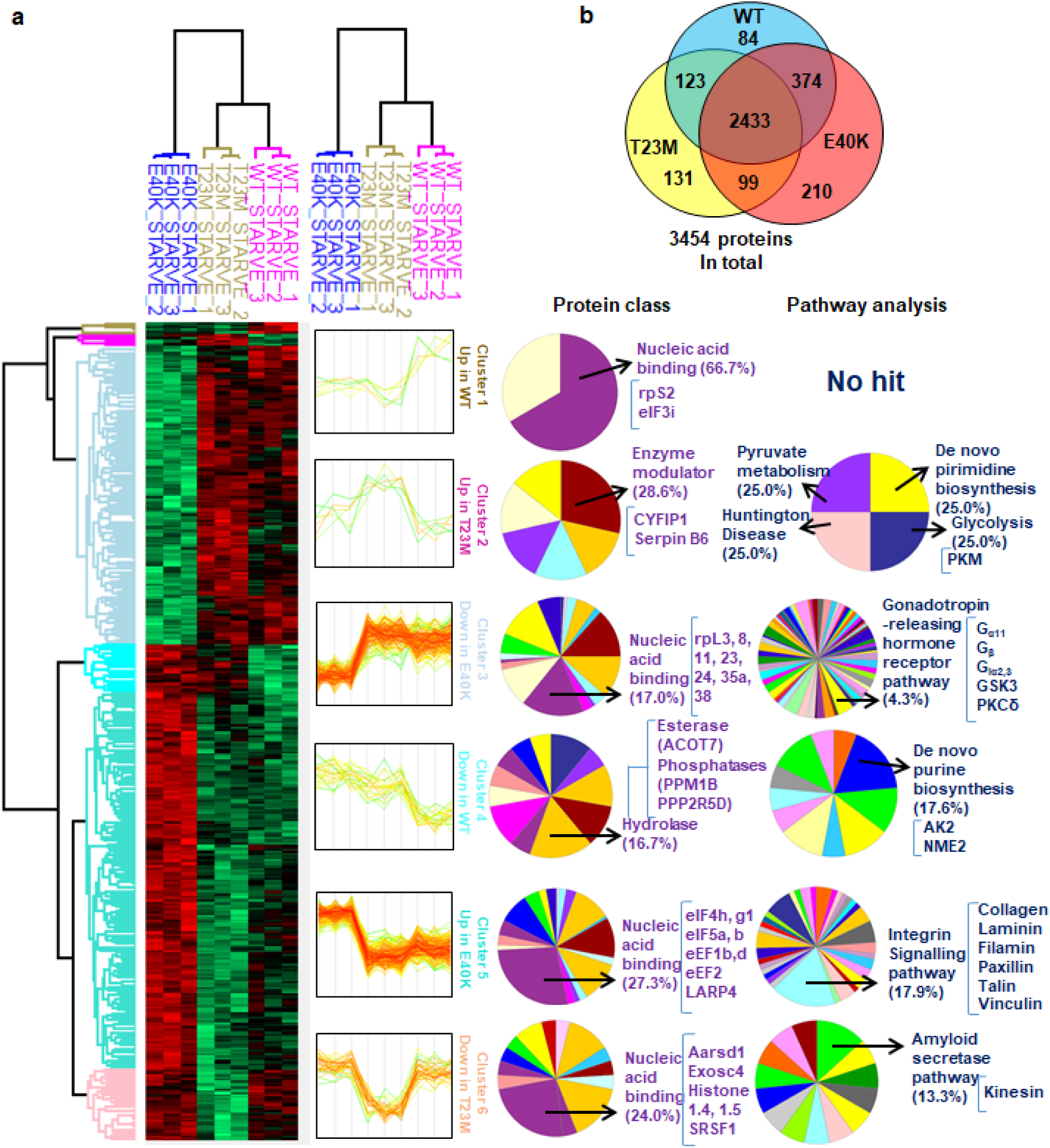

To validate our MS data, we performed western blot analysis for selected proteins, namely PKM2, eEF2, collagen and paxillin. This confirmed that PKM2 was modestly increased only in Rheb-T23M cells and that eEF2 protein is increased in both Rheb-T23M and E40K cells compared to those expressing Rheb-WT (Fig. 6a; quantified Supplementary Fig. S5c). The integrin signalling components paxillin and collagen 2α1 were also increased in Rheb-T23M and E40K cells consistent with the MS data (Supplementary Fig. S5a/b).

**Figure 6.**
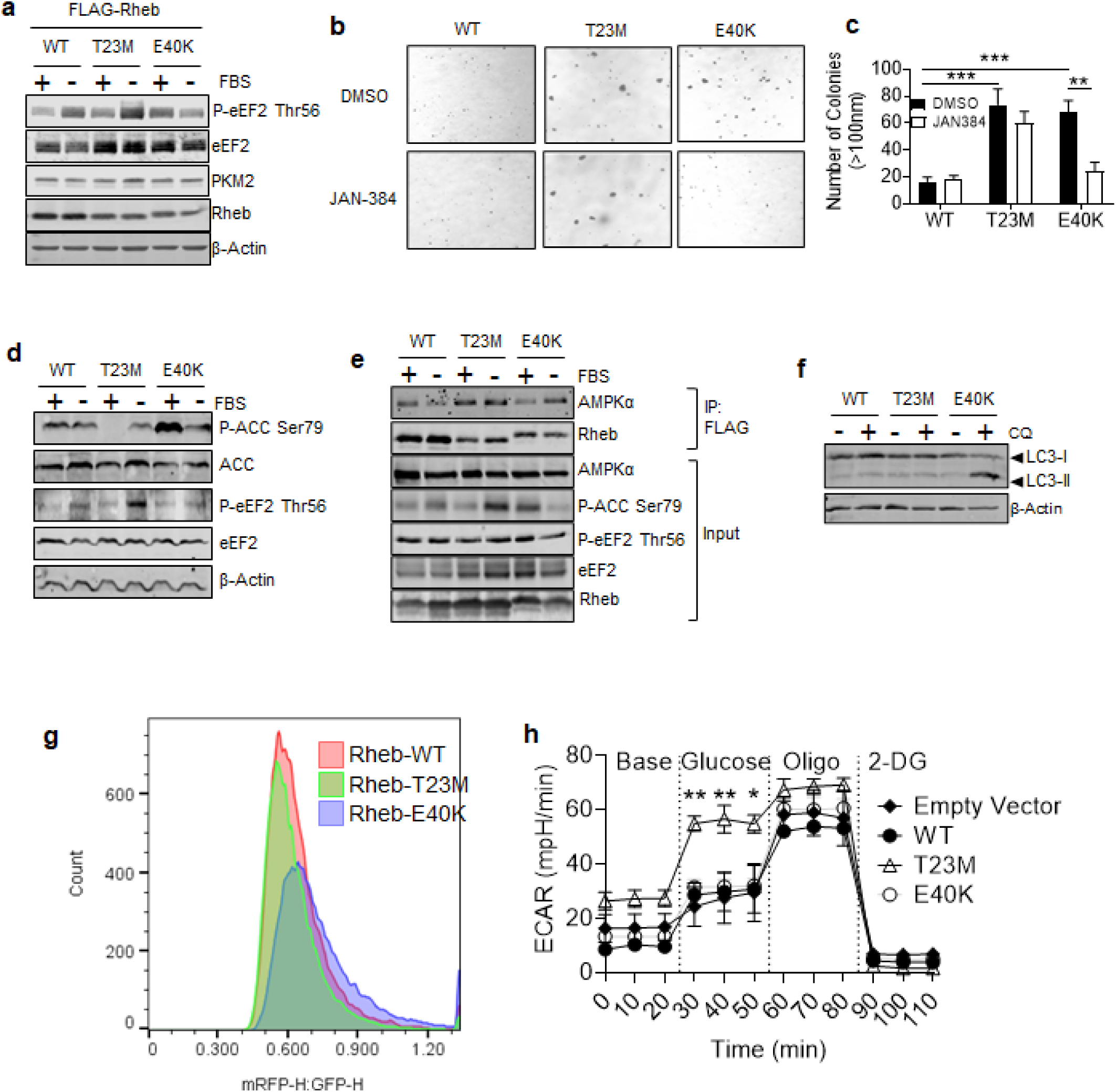

Given that integrin signalling is important for tumour metastasis^34^, we tested whether mutant Rheb proteins affected cells’ ability to migrate or invade using 3D ‘transwell’ assays. Cells expressing Rheb-T23M, Y35N or E40K all showed strikingly higher rates of migration than those transfected with empty vector or expressing Rheb-WT (Supplementary Fig. S6a; quantified Supplementary Fig. S6b). To measure invasion, cells were placed in Transwell chambers where the bottom had been coated with matrigel. Cells expressing the Rheb mutants T23M, Y35N or E40K showed a significantly greater invasive ability than cells expressing Rheb-WT (Supplementary Fig. S6a-c).

### Rheb T23M and E40K differentially regulate AMPK

Interestingly, while total eEF2 protein was increased by both Rheb-T23M and E40K, phosphorylation of eEF2 at Thr56 differed between the mutants with P-eEF2 Thr56 increasing under serum starvation in Rheb-WT and T23M expressing cells but decreasing under the same conditions in Rheb-E40K-expressing cells (Fig. 6a). Under fully supplemented conditions, P-eEF2 was higher in Rheb-E40K cells compared to those expressing Rheb-WT or T23M (Fig. 6a; quantified Supplementary Fig. S5a). Thr56 in eEF2 is only phosphorylated by eEF2K^35^ an atypical protein kinase which has been shown to play important roles in cancer cells ^36^. Given the elevated levels of p-eEF2 in cells expressing Rheb-E40K cells, we hypothesised that eEF2K may be required for tumour growth. In that case, inhibition of eEF2K could result in decreased anchorage-independent growth. As expected for the data presented above (Fig. 3d), in soft agar assays, cells expressing Rheb-T23M or E40K formed large colonies, while treatment with the eEF2K inhibitor JAN-384 prevented growth of cells expressing Rheb-E40K but not Rheb-WT or Rheb-T23M cells, suggesting that phosphorylation of eEF2 is required for the growth under fully-supplemented conditions of cells expressing Rheb-E40K but not of cells expressing Rheb-T23M (Fig. 6b; quantified Fig. 6c).

AMPK is an important upstream activator of eEF2K^37^ whose input overrides the inhibitory effect of mTORC1 on eEF2K activity (the latter being activated by both Rheb mutants, so cannot explain their differing effects on p-eEF2). Given that Rheb-Y35N has previously been shown to regulate AMPK^19^, we hypothesised that AMPK signalling might be differentially regulated by Rheb T23M and E40K. To test this, we starved NIH3T3 cells of serum for 24 h, a condition that activates AMPK (Supplementary Fig. S5d) as judged by the phosphorylation of acetyl-CoA-carboxylase (ACC; at Ser79), which is a well characterised substrate of AMPK and thus an indicator of AMPK activity. P-ACC was strongly increased above its low basal levels under serum-starved conditions in cells expressing Rheb-T23M compared to Rheb-WT, but was strongly decreased in fully supplemented conditions. This suggests that Rheb-T23M may inhibitAMPK under fully supplemented conditions. This is consistent with previous work suggesting that, while Rheb-WT acts as an activator of AMPK, the constitutively-active Rheb mutant Rheb-Y35N inhibits AMPK. Conversely, P-ACC was actually lower in cells expressing Rheb-E40K suggesting AMPK is inhibited under serum starvation conditions in these cells (Fig. 6d). In Rheb immunoprecipitates (using anti-FLAG), we found that AMPK bound less to Rheb-E40K than to Rheb-WT or T23M in cells in fully supplemented medium, but under serum-starved conditions binding of Rheb-E40K to AMPK increased (Fig. 6e; Quantified in Supplementary Figure S5e). These data suggest that while both Rheb-T23M and E40K bind to AMPK, they different roles under different conditions.

To assess further whether increased binding of Rheb-E40K to AMPK is inhibitory under serum-starvation, we investigated their effects on another well characterised process regulated by AMPK, autophagy^38^. Like eEF2K, autophagy is positively regulated by AMPK and negatively regulated by mTORC1. We hypothesised that, since Rheb-E40K binds less to AMPK in fully supplemented medium, AMPK will display higher activity under this conditions in cells expressing this mutant of Rheb. In turn, allowed for activation of AMPK, in, this activated AMPK may drive autophagy even in the face of activated mTORC1 signalling. To test this, we treated NIH3T3 cells stably expressing Rheb-WT, T23M and E40K with the widely-employed inhibitor of autophagy chloroquine (CQ) for 2 h. Cells were then lysed, and western blot analysis performed for LC3. LC3-II is a 17-kDa protein that is recruited to the autophagosomal membrane and is degraded upon fusion with the lysosome. It therefore acts as a marker of autophagic flux since the inhibition of autophagy results in accumulation of LC3-II depending on the rate of autophagy (autophagic flux). LC3-II levels were greatly increased in cells expressing Rheb-E40K that were treated with CQ compared to cells expressing Rheb-WT or T23M. This accumulation of LC3-II is characteristic of increased autophagic flux (Fig. 6f). To further test this, we utilised a previously described^26^ tandem-fluorescent LC3 (tf-LC3) HeLa cell line. As GFP’s fluorescence is quenched at the low intralysosomal pH, an increase in the mRFP1:EGFP ratio indicates LC3 has entered lysosomes (but has not been degraded) pointing to increased autophagic flux. Under fully-supplemented growth conditions, Rheb-E40K caused a modest increase in the mRFP1:EGFP ratio compared to Rheb-WT or T23M (Fig. 6g) indicating that more LC3 is present within the lysosomes of these cells and thus the rate of autophagy is increased. Together, these data suggest that Rheb-T23M and Rheb-E40K may differentially regulate AMPK through binding to AMPK and inhibiting it under distinct conditions, with Rheb-T23M inhibiting AMPK under fully supplemented conditions but not under serum-starvation and Rheb-E40K inhibiting AMPK under serum-starvation but not in fully supplemented conditions, and thus differentially affect autophagy and eEF2K; however future studies are required to fully establish the mechanism by which they differentially affect AMPK.

### Altered metabolism in cells expressing Rheb-T23M

Given that inhibition of eEF2K did not affect the growth of Rheb-T23M-expressing cells in soft agar, we suspected that another pathway(s) might be required for Rheb-T23M to drive cancer. We identified PKM2 levels as being higher in cells expressing Rheb-T23M (Fig. 6a; quantified in Supplementary Fig. S5c/d) suggesting that in these cells anaerobic glycolysis may be favoured over oxidative phosphorylation; this effect [the Warburg effect^39^] has long been known to aid cancer cell proliferation and survival^40^. To test this, we performed a metabolic activity (‘Seahorse®’) assay to measure extracellular acidification rate (ECAR, i.e., due to production of lactate by anaerobic glycolysis).

Cells expressing Rheb-T23M showed much higher basal rates of glycolysis upon addition of the substrate glucose than cells expressing Rheb-WT, E40K or empty-vector (Fig. 6h). As expected, adding oligomycin, which inhibits oxidative phosphorylation by blocking ATP synthase and thus ‘forces’ cells to increase flux through anaerobic glycolysis, markedly enhanced ECAR in Rheb-WT, E40K and empty-vector control cells. In contrast, cells expressing Rheb-T23M showed only a modest further increase in their already elevated ECAR. Maximal rates of ECAR (‘glycolytic capacity’) were similar in all cases. These data indicate that Rheb-T23M drives basal glycolytic rates much closer to the cells’ glycolytic capacity than is normally the case; thus, cells expressing Rheb-T23M appear to favour anaerobic glycolysis over oxidative mitochondrial metabolism, likely reflecting their increased levels of PKM2. This likely also contributes to their enhanced oncogenic potential.

## Discussion

It is now clear that signalling through mTORC1 is deregulated in many cancers, due to mutation or loss of components of oncogenic upstream signalling pathways that impinge on TSC and thus Rheb^41^, such as proteins involved in the classical MAP kinase (ERK) and PI 3-kinase/PKB (Akt) pathways including Ras or Pi 3-kinase, which are mutated in many cancers. Here we show that multiple mutations in the Rheb-GTPase, a proximal regulator of mTORC1^8, 42-45^, that have been identified in genomic studies of patient-derived tumours but previously considered to be background mutations or genetic noise^19^, also lead to constitutive activity and promote signalling through mTORC1. Our results show that, in addition to the previously-reported Rheb-Y35N mutant^19, 20, 28, 30^, several further mutants identified in multiple cancer genome studies on a variety of different cancers are resistant to negative control by TSC1/2 and thus promote aberrantly active mTORC1 signalling.

Consistent with previous studies^28^, we found that the Rheb mutants S21L, T23M, G29S, Y35N and E40K, are partially or wholly resistant to the GAP activity of TSC2, both when TSC1/2 is overexpressed in cells as well an in an *in vitro* GTPase assay. These data therefore suggest that these Rheb mutants do not drive constitutive mTORC1 signalling through Rheb-mediated MAPK activation^19, 20^, but rather through reduced sensitivity to the GAP activity of TSC2. Recent studies have also reported that Rheb-Y35N causes constitutive activation of MAPK leading to dysregulated mTORC1 signalling through continual inhibition of TSC2^19, 20^. We show that Rheb-T23M, G29S, Y35N and E40K also increase P-Erk1/2 in fully supplemented medium (Supplementary Fig. S1C), but that neither pharmacological inhibition nor upstream inhibition of MAPK prevents the stimulation of mTORC1 signalling by Rheb mutants. Thus, the ability of these mutants to stimulate mTORC1 signalling does not require the MEK/ERK pathway.

Combined data from three cancer genomics databases (COSMIC, The Cancer Genome Atlas and the Broad Institute Cancer Cell Line Encyclopedia) revealed that Rheb mutations occur in some cancers with the most common ones being, in descending order, Y35N, Q57*, T23M and E40K. The frequency of Rheb-Q57* seems perhaps surprising as this truncation removes several important structural and functional domains, most notably the Switch II region (residues 63-79) which is crucial for Rheb function. It is therefore unlikely that Rheb-Q57* is constitutively active or even functional. Given that NIH3T3 cells stably expressing Rheb-T23M and E40K showed faster proliferation and robust growth in soft agar as well as increased growth in an in vivo syngeneic tumour graft, the Rheb-T23M and E40K variants are not simply background (‘passenger’) mutations, but instead likely act as strong driver oncogenes.

To assess global changes in protein expression brought about by Rheb-driven, constitutively-active mTORC1 signalling, we performed MS analysis on NIH3T3 cells stably expressing Rheb-WT, T23M or E40K grown in DMEM lacking FBS. As our data imply that both mutants drive constitutive mTORC1 signalling through the same mechanism (insensitivity to TSC1/2), we were surprised to observe other quite distinct changes in the proteome suggesting that Rheb, or at least these Rheb mutants, may exert functions additional activating mTORC1 signalling. Given that E40 is located within the Switch II region of Rheb, which has been shown to be important for mTORC1 binding^46^, it is possible this mutation alters the interaction between Rheb and other binding partners. In contrast, since T23 is located adjacent to, but not within, the P-loop, the T23M mutation likely affects only GTP hydrolysis and would be expected only to maintain Rheb activity without significantly altering protein-protein interactions. This notion of differential effects of the two mutations is supported by our observation that the interaction of Rheb T23M and E40K with AMPK, a known binding partner of Rheb^19, 47^, responds differently to serum starvation. The crystal structures of Rheb-T23M and E40K would need to be resolved in order to determine whether these mutations do indeed differentially alter Rheb-protein interactions.

We show that, as well as promoting constitutive mTORC1 activation, Rheb-T23M and E40K each regulate additional pro-oncogenic pathways that may drive cancer. We show that eEF2K activity is required for anchorage-independent growth of NIH3T3 cells expressing Rheb-E40K, suggesting that constitutive mTORC1 activation alone is insufficient to drive such growth in this context and that eEF2K activity is also required. Why eEF2K should be necessary to drive cancer growth in cells expressing Rheb-E40K but not Rheb-T23M initially appears puzzling. It has previously been shown that TSC1/2 null cells, which mimic the phenotype of constitutively active Rheb mutants, expend more ATP and become reliant on glucose for survival due to the increase in protein synthesis that accompanies mTORC1 activation^48^. We have also previously shown that AMPK phosphorylates eEF2K in response to poor intracellular energy status^37^. It is possible that the pro-survival effects of AMPK complement the pro-growth effects of mTORC1, thus producing the highly advantageous conditions for sustained cancer growth. This would be further aided by the AMPK mediated activation of autophagy, which in this context appears to over-ride the inhibitory effects of mTORC1, as it would help provide the necessary macronutrients to sustain growth. The ability to drive both mTORC1 signalling and autophagy would be a significant advantage to cancer cells especially in poor nutritional conditions where autophagy can provide precursors for the macromolecular synthesis pathways that are driven by mTORC1. This hypothesis also offers an explanation as to why eEF2K is not required for growth of cells expressing Rheb-T23M. These cells display a preference for utilising anaerobic glycolysis linked to the upregulation of PKM2. This effect, commonly called the Warburg effect, is a well-known feature of tumour cells and allows for the rapid production of ATP. In this context, this may act as a compensatory mechanism to provide cells with the energy required for sustained growth without the activation of a pro-survival pathway such as AMPK. Future studies are required to test this hypothesis and to understand both the biological relevance of this observed difference between Rheb-T23M and E40K as well as the mechanisms responsible.

In summary, we show here that several tumour-associated mutations in Rheb are constitutively active and act as driver oncogenes. We show that they drive constitutive mTORC1 signalling through insensitivity to TSC2’s GAP activity (and not through constitutive activation of MAPK as previously suggested for another Rheb mutant). Finally, our data reveal that Rheb-T23M and E40K regulate distinct, cancer-associated pathways. These findings also suggest that unique ‘bespoke’ combination therapies may be utilised to treat cancers harbouring different Rheb mutants.

## Conflicts of Interest

The Authors declare no potential conflicts of interest

## Notes

Financial Support: This work was initiated with funding from the UK Biotechnology and Biological Sciences Research Council (to CGP) and then supported by SAHMRI. SDP acknowledges support by a Scholarship from the Australian Government Research Training Program (RTP). JX acknowledges a SAHMRI early/mid-career seed funding grant.

### Competing Interest Statement

The authors have declared no competing interest.

