## Supplementary figures 1-6 for "TSC-Insensitive Rheb Mutations Induce Oncogenic Transformation Through a Combination of Hyperactive mTORC1 Signalling and Metabolic Reprogramming"

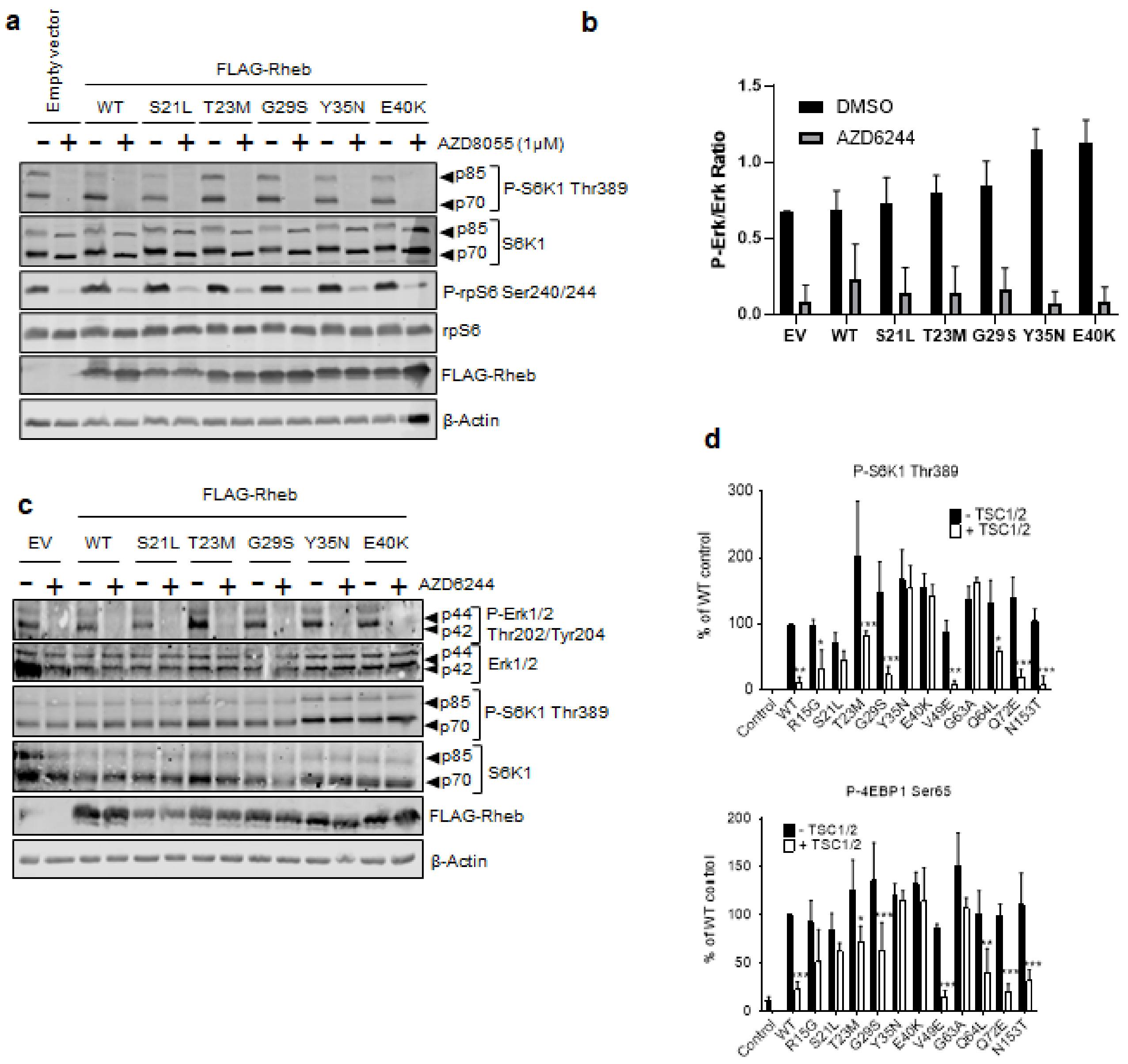


**Supplementary Figure S1**

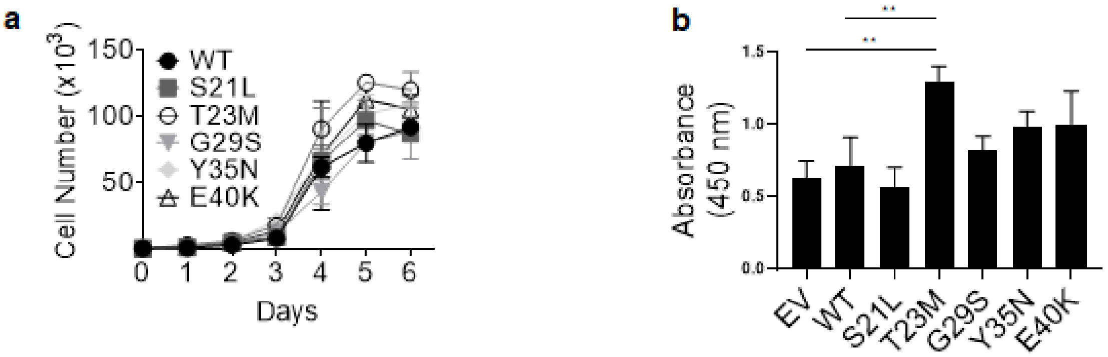


**Supplementary Figure S2**

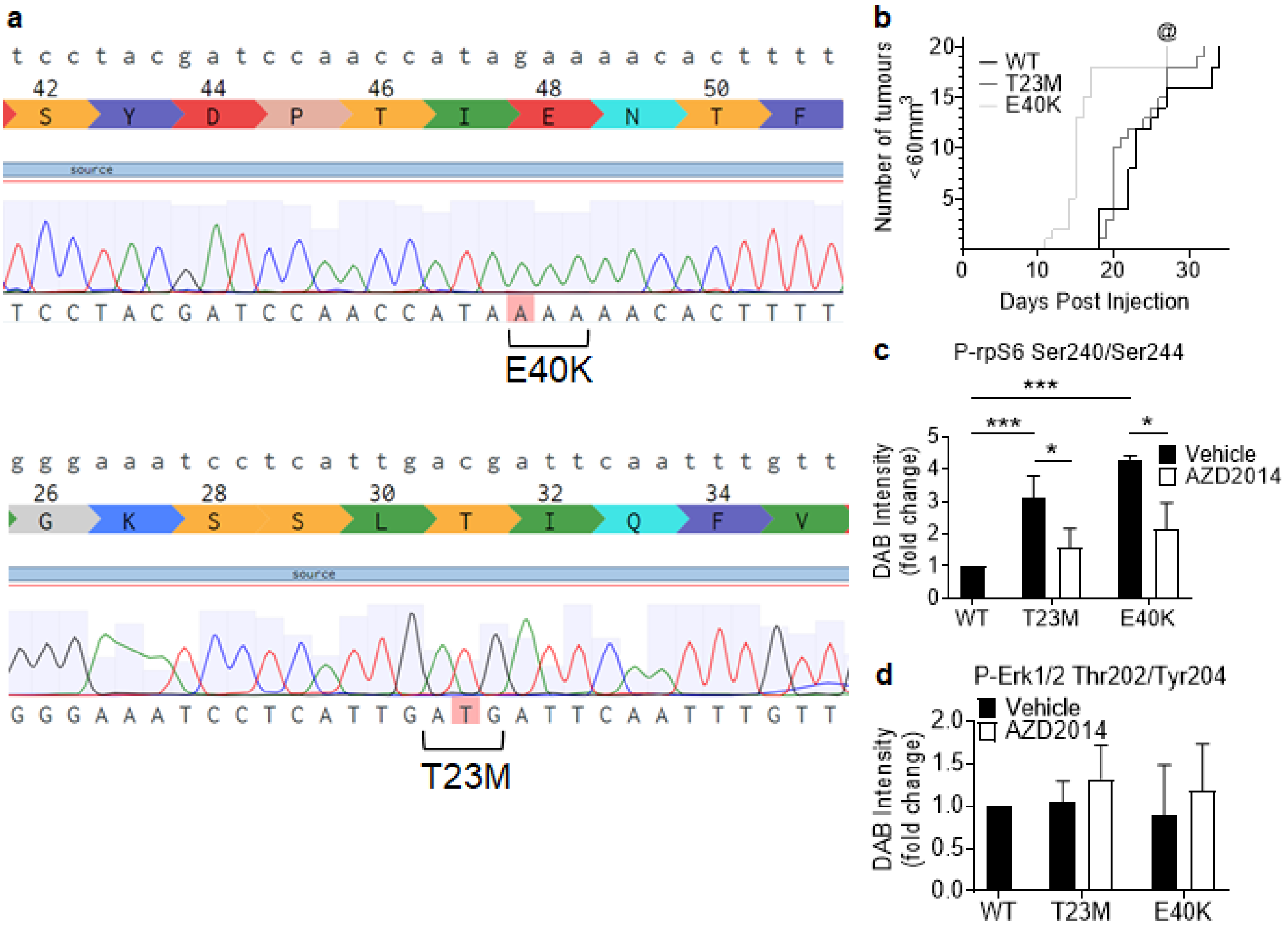


**Supplementary Figure S3**

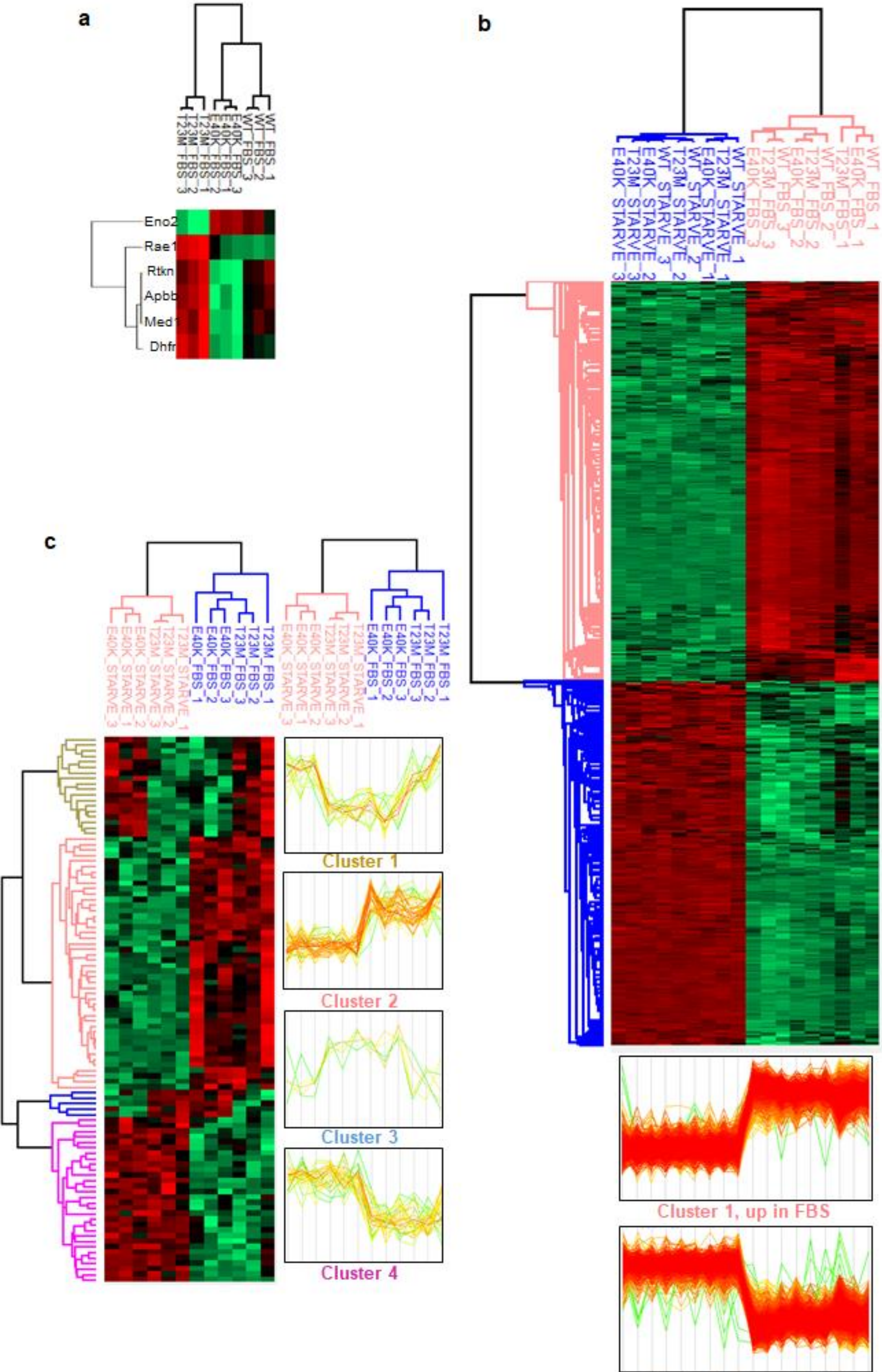


**Supplementary Figure S4**

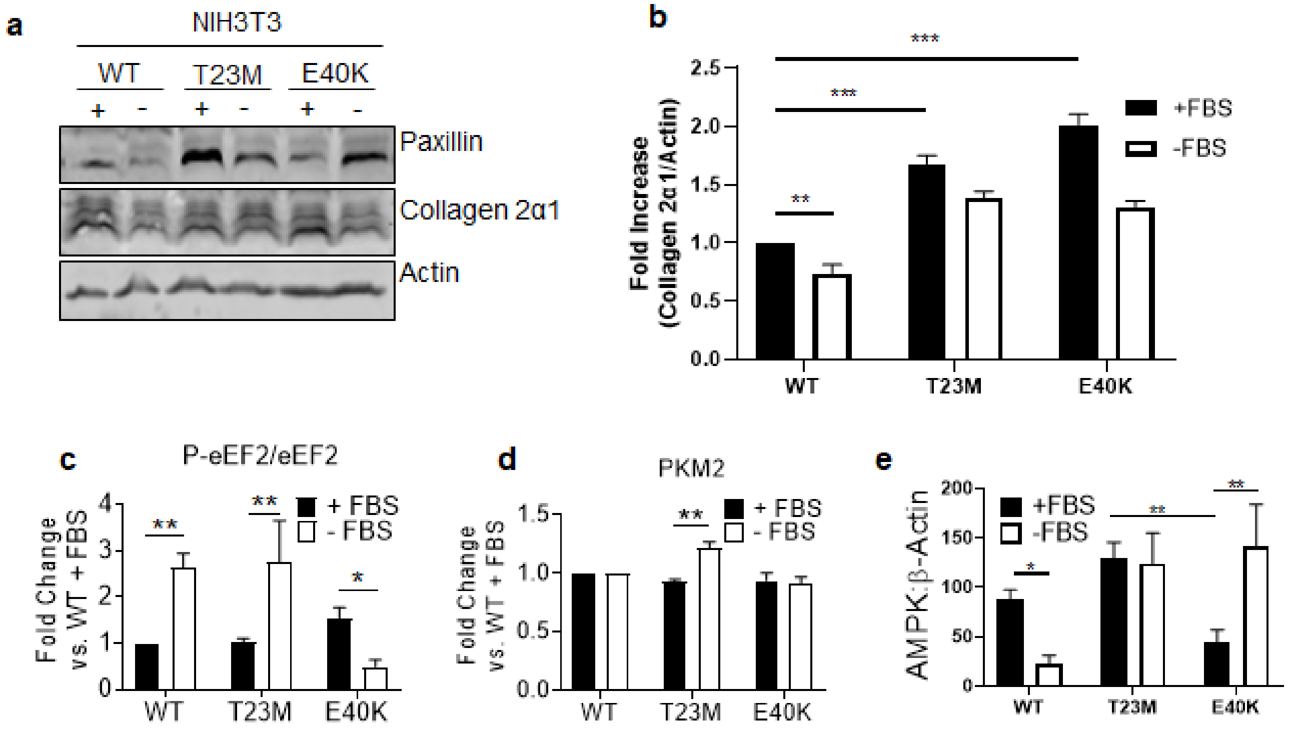


**Supplementary Figure S5**

a

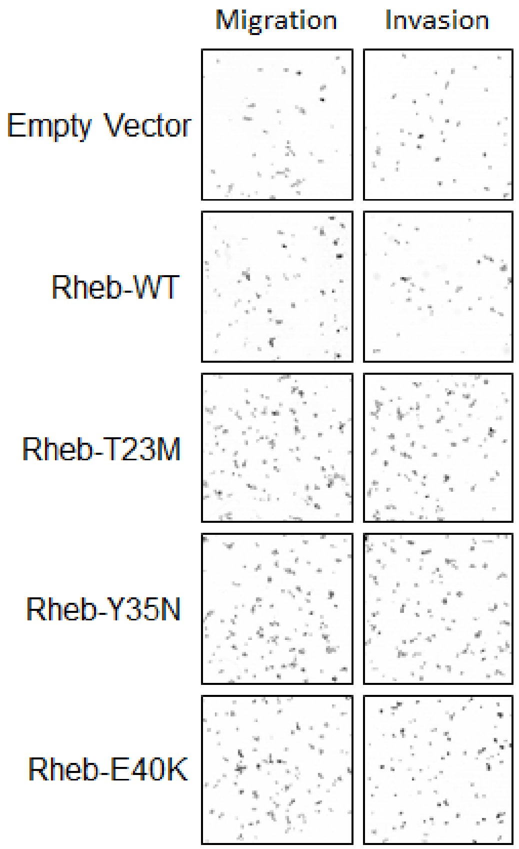


b

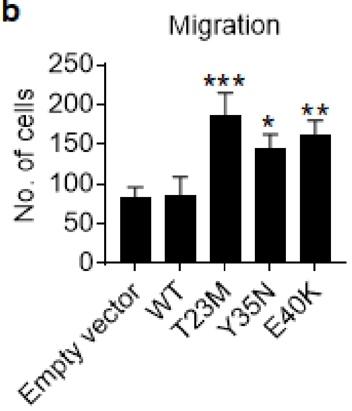


c

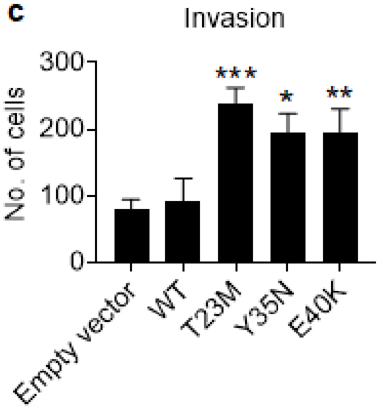


d

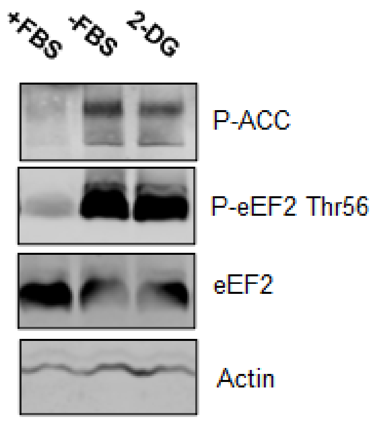


**Supplementary Figure S6**
